# Stage-specific, morphological and molecular markers of encystation in *Giardia lamblia*

**DOI:** 10.1101/2021.02.01.429238

**Authors:** Elizabeth B. Thomas, Renaldo Sutanto, Richard S. Johnson, Han-Wei Shih, Jana Krtková, Michael J. MacCoss, Alexander R. Paredez

## Abstract

Differentiation into environmentally resistant cysts is required for transmission of the ubiquitous intestinal parasite *Giardia lamblia*. Encystation in *Giardia* requires the production, processing and transport of Cyst Wall Proteins (CWPs) in developmentally-induced, Golgi-like, Encystation Specific Vesicles (ESVs). Progress through this trafficking pathway can be followed by tracking CWP localization over time. However, there is no recognized system to distinguish the advancing stages of this process which can complete at variable rates depending how encystation is induced. Here we propose a staging system for encysting *Giardia* based on the morphology of CWP1-stained ESVs. We demonstrate the molecular distinctiveness of maturing ESVs at these stages by following *Gl*Rab GTPases through encystation. Previously, we established that *Giardia*’s sole Rho family GTPase, *Gl*Rac, associates with ESVs and has a role in regulating their maturation and the secretion of their cargo. As a proof of principle, we delineate the relationship between *Gl*Rac and ESV stages. Through proteomic studies, we identify putative interactors of *Gl*Rac that could be used as additional stage-specific ESV markers. This staging system provides a common descriptor of ESV maturation regardless of the source of encysting cells. Furthermore, the identified set of molecular markers for ESV stages will be a powerful tool for characterizing trafficking mutants that impair ESV maturation and morphology.

**Importance:** *Giardiasis* is a diarrheal disease that affects 280 million people worldwide. It is caused by *Giardia lamblia*, a protozoan parasite which rely on differentiating from host-dwelling trophozoites to environmentally-resistant cysts for transmission and survival. This encystation process requires the transport of Cyst Wall Proteins (1-3) within membrane-bound compartments called Encystation Specific Vesicles (ESV) from the endoplasmic reticulum to the surface of the cell. The whole process takes 24 hours to complete and these compartments are the only recognizable equivalent of Golgi apparatus in this minimalistic organism. Progress of this trafficking pathway can be followed by localizing Cyst Wall Protein 1 over time post induction of encystation but this can be ambiguous when specific molecular events need to be specified. Here we propose a staging system that is based on ESV morphology changes by capitalizing on the secretory/processing events we already know they represent. We validate the molecular distinctiveness of these stages by following *Giardia* Rabs through the pathway and characterize putative interactors of an established regulator of encystation, *Gl*Rac, to provide additional stage-specific molecular markers. This staging system will provide a definitive, yet adaptable, framework to map out functions of yet-to-be discovered players of this important pathway.

## Introduction

*Giardia lamblia* (syn. *Giardia intestinalis* and *Giardia duodenalis*) is a major intestinal parasite which infects more than 280 million people every year (Lane & Lloyd, 2002). The lifecycle of this diplomonad protozoan is simple, featuring only two stages – the binucleate, double-diploid, proliferative trophozoites which non-invasively colonizes host intestines and the environmentally-resistant, infectious, non-motile cysts that are shed in host’s feces. Regulation of encystation ensures the production of viable cysts and promotes transmission of this ubiquitous parasite. Being a popular life-cycle strategy also adopted by other protozoan parasites, studying this differentiation process is important and *Giardia* is the best-developed model available (Eichinger, 2001).

*Giardia* encystation requires the construction of its protective cyst wall, an extracellular matrix composed of Cyst Wall Material (CWM). CWM contains three paralogous Cyst Wall Proteins (CWP1-3) and a unique β-1,3-linked N-acetylgalactosamine (GalNAc) homopolymer (Gerwig et al., 2002; Lujan et al., 1995; Sun et al., 2003). When induced, large quantities of CWPs are synthesized and transported from the endoplasmic reticulum (ER) to the cell surface in membrane-bound organelles called Encystation-Specific Vesicles (ESVs). *Giardia* lacks classical Golgi apparatus. However, since nascent ESVs arise from ER-exit sites (ERES; Faso et al. 2013) and are marked by several Golgi markers they are thought to be developmentally-induced Golgi (Marti et al., 2003). This view is supported by the roles ESVs play as the only recognizable post-ER delay compartments. ESVs feature machinery needed for the post-translational processing and subsequent partitioning of CWPs into distinct phases (Davids et al., 2004; Reiner et al., 2001; Slavin et al., 2002). After proteolytic cleavage of CWP2, CWP1 and the N-terminal end of cleaved CWP2 are sorted into the outer fluid phase while CWP3 and C-terminal end of cleaved CWP2 remain as the inner condensed core (DuBois et al., 2008; Konrad et al., 2010; María C. Touz et al., 2002). Additionally, ESVs coordinate secretion of CWM to the cell surface, likely mediated by a higher order networking structure (Štefanić et al., 2009). Processed CWPs are deposited sequentially; the fluid phase first at a rapid rate where binding of CWP1 to the cell surface is mediated by its lectin binding domain that recognizes GalNac fibrils on the surface of the encysting cell (Chatterjee et al., 2010). This is followed by slower secretion of the condensed phase (Konrad et al., 2010; Štefanić et al., 2009). These events are trackable by following ESV morphology and CWP localization.

As a lab-inducible and -tractable secretory pathway of a minimalistic organism, *Giardia*’s encystation process provides a unique opportunity to uncover the constraining principles of membrane trafficking. Despite fundamental differences in compartment organization, canonical membrane trafficking players continue to perform conserved roles in *Giardia* (Maria C. Touz & Zamponi, 2017). The accumulation of CWP and *de novo* ESV biogenesis at the ERES is dependent on COPII and the small GTPase, SarI – vesicle coat proteins that transport cargo from the rough ER to the Golgi apparatus in higher eukaryotes (Štefanić et al., 2009). Additionally, Arf1, a small GTPase that canonically plays a central role in intra-Golgi transport by regulating COPI and clathrin membrane coats, is required for ESV maturation; inhibiting Arf1 activity interfered with the transport and secretion of CWM to the cell surface (Štefanić et al., 2009). The sole Rho GTPase in *Giardia, Gl*Rac, whose homologs are known to coordinate vesicle trafficking and the cytoskeleton in plants and animals, was found to regulate CWP trafficking and secretion (Krtková et al., 2016).

Similar to its homologs, *Gl*Rac is thought to regulate *Giardia* encystation by acting as a molecular switch for the recruitment and regulation of effector proteins which drive the progress of encystation. We previously demonstrated that *Gl*Rac is required for the temporal coordination of CWP1 production, ESV maturation, and CWP1 secretion (Krtková et al., 2016). The relationship between *Gl*Rac and its roles in CWP trafficking are complicated by the dynamic association of *Gl*Rac with ESVs (Krtková et al., 2016). While it is recognized that ESVs go through stages of maturation (Konrad et al., 2010), there is no established criteria for specifically identifying these stages. Additionally, timing post induction of encystation has been used to stage ESVs, but the amount of time it takes to encyst varies by the method used to promote encystation (Einarsson et al., 2016; Konrad et al., 2010; Luján et al., 1996). For a direct comparison of CWP1 and CWP2 induction using these methods see Fig. S1. Our efforts to specify the molecular events that coincided with *Gl*Rac activity at ESVs, highlighted the need for a standardized system to distinguish the advancing stages of encystation. Here, we develop a system for staging encysting *Giardia* cells that is based on our current understanding of molecular events during encystation and observed by changes in ESV morphology (Konrad et al., 2010). Furthermore, this would allow us to navigate around ambiguities introduced by variation in the timing and efficiency of encystation when different protocols for induction are used. Our survey of *Giardia* Rabs, small GTPases that are known to control the specificity and directionality of membrane trafficking pathways and mark specific organelles, highlight the molecular distinctiveness between the stages we propose. As a proof of principle, we characterized putative interactors of *Gl*Rac as identified by proteomics and confirm their association with ESVs. This work would facilitate future studies where the functions of *Gl*Rac effectors would be precisely mapped in this crucial trafficking pathway.

## Materials and Methods

### Strain and culture conditions

*G. lamblia* strain WB clone C6 (ATCC 50803; American Type Culture Collection) was cultured in TYDK medium (per 100ml; 2 g casein digest, 1 g yeast extract, 1 g glucose, 0.2 g NaCl, 0.06 g KH_2_PO_4_, 0.095 g K_2_HPO_4_, 0.2 g L-cysteine, 0.02 g L-ascorbic acid, 1.2 mg ferric ammonium citrate) supplemented with 10% adult bovine serum with pH adjusted to 7.1. To induce encystation in trophozoites, the two-step encystation protocol was followed. Trophozoite cultures were first grown to confluency (∼1 × 10^6^ cells/ml) for 36 h in pre-encystation medium (same as growth medium above but without bile and pH at 6.8). Encystation was induced by switching culture medium to encystation medium (same as growth medium but pH 7.8 and supplemented with 10 g/l bovine/ovine bile instead of bovine bile) and incubating the cells further for either 8 h or 24 has indicated.

### Vector construction

All constructs were made using traditional cloning, Gibson assembly (Gibson et al., 2009) or In-Fusion kit (Takara Bio); for full details and Gene Accession numbers, consult Table S1. Note, constructs made to tag *Giardia* Rabs and putative *Gl*Rac-interactors were designed for stable integration by homologous recombination into the *Giardia* genome for endogenous expression. Generation of the native promoter morpholino-sensitive (ms) HALO_C18_*Gl*Rac_PuroR (designed for endogenous expression) and PAK promoter_CRIB_N11_mNG_3HA_NeoR (designed for episomal expression) constructs have been described previously (Hardin et al., 2021, submitted). Plasmid backbones used to build constructs in this paper were sourced from Gourguechon & Cande, 2011, Krtková et al., 2016, Michaels et al., 2020 and Paredez et al., 2014.

For transfection, 5 to 50 µg of DNA was electroporated (375 V, 1,000 µF, 750 Ω; GenePulser Xcell; BioRad, Hercules, CA) into trophozoites. Following electroporation, the cells were added to 13 ml pre-warmed, fresh TYDK medium and allowed to recover at 37°C overnight before beginning selection with G418 or Puromycin for 4-7 days. Strains were maintained at a final concentration of 30 μg/ml G418 or 0.3 mg/ml Neomycin.

### Luciferase assay

Confluent cell cultures were incubated with the three different encystation media (two-step, Uppsala, and lipoprotein-deficient) for 8 h and 20 h, then pelleted and resuspended with Hepes-Buffered Saline. 200 µl diluted cells (2 x10^4^ cells) and 50 µl (8 mg/ml) D-luciferin (GoldBio, USA) were loaded into each well. Plates were incubated for time increment from 5 min to 30 min at 37°C. Luciferase activity was determined with an Envision MultiLabel Plate Reader (Perkin Elmer, USA).

### Immunofluorescence microscopy

Immunofluorescence assays were performed as described previously (Krtková et al., 2016), To detect 3HA tag, anti-HA rat monoclonal antibodies 3F10 (Roche) diluted to 1:125 followed by Alexa 488-conjugated anti-rat antibody (Molecular probes) diluted to 1:250 were used. To detect HALO tag 0.5µM Janelia Fluor 549 (Promega) dye or HaloTag® TMR Ligand (Promega) were used. CWP1 was detected with Alexa 647-conjugated anti-CWP1 antibody (Waterborne, New Orleans, LA)

Fluorescent images were acquired on DeltaVision Elite microscope using a 100×, 1.4-numerical aperture objective and a PCO Edge sCMOS camera. Deconvolution was performed with SoftWorx (API, Issaquah, WA) and images were analyzed using Fiji, ImageJ (Schindelin et al., 2012). Pearson Coeffecient analysis and Costes’ automatic thresholding were obtained using the JACoP plugin for ImageJ (Bolte & Cordelieres, 2006). 3D viewing and manual scoring of cells were performed using Imaris (Bitplane, version 8.9) Figures were assembled using Adobe Photoshop.

### Affinity purification

Affinity purification of OneSTrEP-*Gl*Rac was done according to a previously used protocol (Paredez et al., 2014) Note that two-step encystation protocol was followed during culturing for the 8 h p.i.e. experiments. The resulting elutes were analyzed with Liquid Chromatography with tandem mass spectrometry.

### Mass Spectrometry

Samples were prepared using the FASP method (Wiśniewski et al., 2009). Briefly, the samples were concentrated in an Amicon Ultra 10K filter, washed twice with 25 mM ammonium bicarbonate (ABC), and the disulfides reduced using 10 mM tris 2-carboxyethyl phosphine (TCEP) for 1 h at 37°C. The resulting thiols were alkylated with 12 mM iodoacetamide for 20 min prior to spinning out the liquid. The proteins were washed twice with 100 µl 50 mM ABC. Digestion occurred after addition of 1 µg trypsin (Promega sequencing grade) in 200 µl 50 mM ABC, and overnight incubation at 37°C. The peptides were spun out of the filter and dried using a vacuum centrifuge.

All mass spectrometry was performed on a Velos Pro (Thermo) with an EasyLC HPLC and autosampler (Thermo). The dried pulldowns were solubilized in 25 µl of loading buffer (0.1% trifluoroacetic acid and 2% acetonitrile in water), and 6 µl was injected via the autosampler onto a 150-μm Kasil fritted trap packed with Reprosil-Pur C18-AQ (3-μm bead diameter, Dr. Maisch) to a bed length of 2 cm at a flow rate of 2 µl/min. After loading and desalting using a total volume of 8 µl of loading buffer, the trap was brought on-line with a pulled fused-silica capillary tip (75-μm i.d.) packed to a length of 25 cm with the same Dr. Maisch beads. Peptides were eluted off the column using a gradient of 2-35% acetonitrile in 0.1% formic acid over 120 min, followed by 35-60% acetonitrile over 5 min at a flow rate of 250 nl/min. The mass spectrometer was operated using electrospray ionization (2 kV) with the heated transfer tube at 25°C using data dependent acquisition (DDA), whereby a mass spectrum (m/z 400-1600, normal scan rate) was acquired with up to 15 MS/MS spectra (rapid scan rate) of the most intense precursors found in the MS1 scan.

Tandem mass spectra were searched against the protein sequence database that was downloaded from GiardiaDB, using the computer program Comet (Eng et al., 2013) Iodoacetamide was a fixed modification of cysteine, and oxidized methionine was treated as a variable modification. Precursor mass tolerance was 2 Da, and fragment ion tolerance was +/-0.5 Da. Discrimination of correct and decoy spectra was performed using Percolator (Käll et al., 2007) with a 1% q-value cutoff. Proteins that had more than one unique peptide and significantly higher values for normalized total spectral counts (as determined by Fisher Exact Test with a Bonferroni multiple testing correction with an alpha of 0.01) in the OneSTrEP-*Gl*Rac pulldown sample compared to its counterpart WT control sample, were noted to be hits.

## Results

### Encystation Staging

Encystation was induced with a two-step protocol where *Giardia* trophozoites were initially cultured in no-bile, low-pH media (pH 6.8) and then moved into high-bile, high pH media (pH 7.8) (Boucher & Gillin, 1990). This is thought to maximally synchronize the encystation process (Konrad et al., 2010). In our hands, while some level of synchronization could be achieved from this method, cells remained in various stages within the process of cyst development, as judged by localizing CWP1 and visualizing ESV morphology (Fig. 1). We, therefore, sought to increase the resolution of staging encysting cells by going beyond noting hours post induction of encystation (h p.i.e.) and categorizing each cell based on the morphology of CWP1-stained ESVs instead (Fig. 1).

**Fig. 1.**
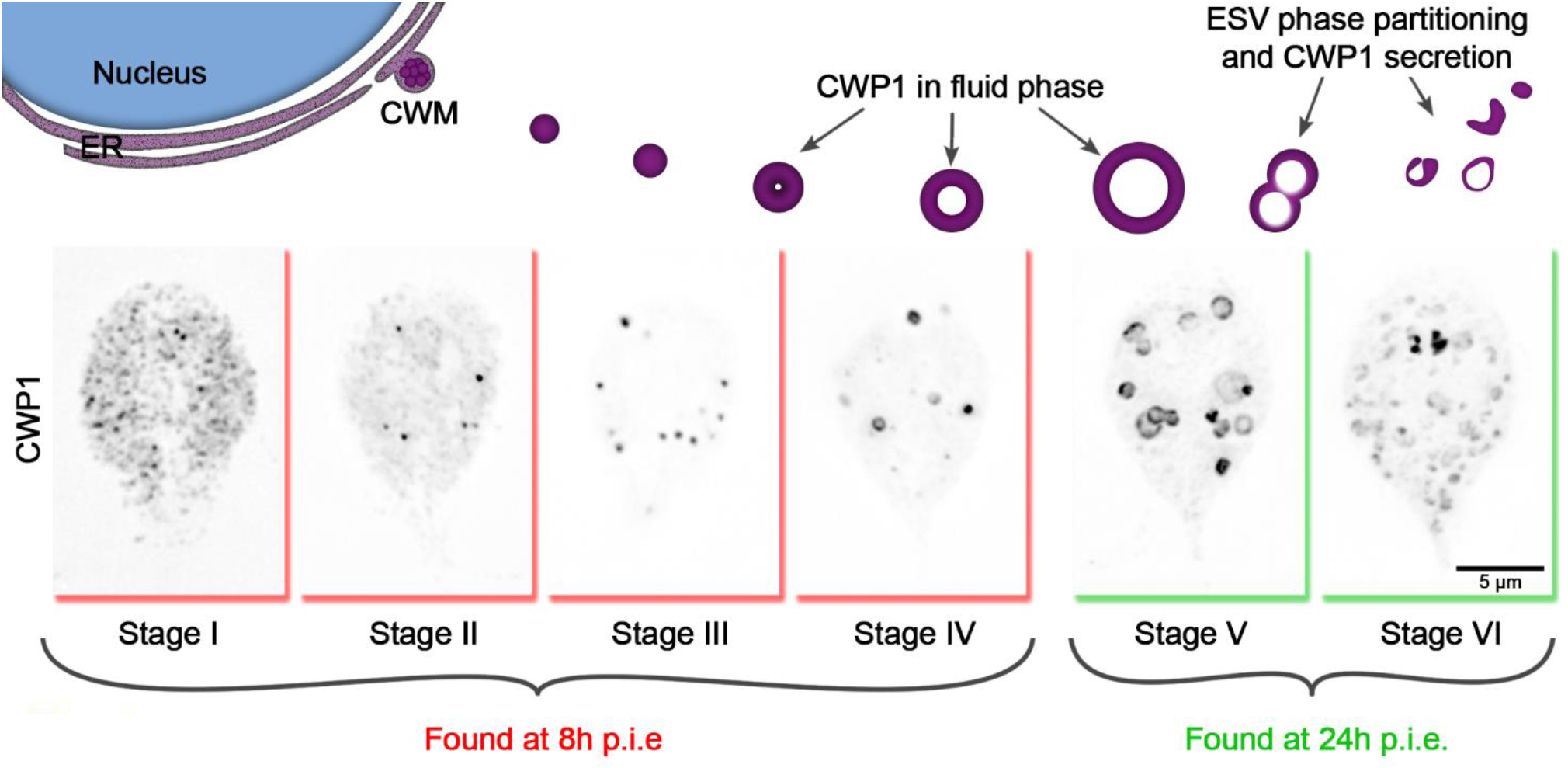
Cyst wall protein trafficking in ESVs during encystation can be divided into stages. Graphic showing trafficking of CWP1-containing ESVs. Once *Giardia* cells sense the signal to encyst, large amounts of CWM are produced in the ER (Stage I), which then accumulate at ER exit sites (Stage II) and are secreted out into the cytoplasm compartmentalized in ESVs (Stage III). As CWP cargo are sorted and processed, they are separated into phases with CWP1 now being confined to the outer fluid-phase of the ESVs (Stage IV) which grow larger in size as they continue to mature (Stage V). The distinct phases of ESVs are then partitioned into separate vesicles (Stage VI) which are subsequently sequentially secreted to form *Giardia’s* cyst wall. These stages can be tracked by visualizing CWP1 and tracking ESV morphology. Cells shown here were harvested and fixed at 8h p.i.e. and 24h p.i.e., then stained with CWP1 antibody.

Based on our current understanding of the sequence of events involved in encystation, we propose the following key for staging encysting cells (Fig. 1): Stage I –CWP1 localizes to the ER; Stage II – CWP1 localize in the ER and also in ER-associated punctate structures thought to be ER-exit sites (Faso et al., 2013); Stage III – CWP1 localizes homogenously in small ESVs; Stage IV – CWP1 localize to doughnut-shaped structures as a result of CWP2 being proteolytically processed to drive core condensation which pushes fluid phase CWP1 to the vesicle periphery. Stage V – found in cultures induced to encyst longer than 8 h (24 h p.i.e in this study); CWP1 now localizes to large doughnut-shaped structures of matured ESVs about to be partitioned from condensed-phase cyst wall material (CWM) into separate vesicles. Stage VI – CWP1 is present in smaller separate vesicles that are close to the surface of the cells and ready to be secreted out to form the cyst wall. Our focus here is on ESV maturation therefore we did not analyze partially to fully formed cysts. Nonetheless, these staging parameters can be expanded to include them in the future.

### *Giardia* Rab GTPases associate to ESVs in stage-specific manner

It is well established that different compartments of the membrane trafficking pathway feature unique molecular identities. Rab GTPases, through the recruitment of specific effectors, regulate the trafficking of each other through the endocytic/sorting/secretory pathway (Pfeffer, 2017). The resulting spatio-temporal specific recruitment of Rabs help direct cargo traffic. Rab GTPases can, therefore, be used as identity markers for differentiating trafficking compartments. We hypothesized that a subset of the nine Rab GTPases encoded in the *Giardia* genome would also demarcate ESVs as they mature, providing unique molecular markers for different stages of these compartments as they sort and secrete CWPs. Each *Gl*Rab was tagged on the N-terminus with mNeonGreen (mNG) to visualize their localization and the cells were subjected to the two-step encystation process before being processed for immunofluorescence assays. Cells were staged as described above and the degree of colocalization between CWP1 and the *Gl*Rab GTPases were determined manually.

To cross-check our qualitative scoring, colocalization was verified by calculating Pearson Coeffecients of 3 cells per stage following Costes’ automatic thresholding; note that these global statistical measurements are not effective for highlighting qualitative differences when proteins display complex localization patterns (Bolte & Cordelières, 2006; Fig. S2). Therefore, we manually scored 15 to 20 cells per stage; after viewing images in 3D, they were given a score between 0-5, with higher scores being granted to cells featuring more robust recruitment of tagged-Rab proteins colocalizing with CWP1-stained vesicles (Fig. S3). Consistency between scores assigned by two team members for a sample subset of cells gave us confidence that this approach is reproducible. Of interest, were Rabs with CWP1 colocalization scores that peaked at the different encystation stages – *Gl*Rab2a at Stages I and II, *Gl*RabA at Stage III, *Gl*Rab D at Stage IV and *Gl*Rab1a at Stages V and VI (Fig. 2). *Gl*Rab32 did not localize to ESVs at any of the stages we looked at (Fig. S3). Multiple attempts at tagging *Gl*Rab2b were unsuccessful and therefore this Rab GTPase was not included in our analysis. Our data is consistent with the cisternal maturation model where ESVs cargo remains in place and the molecular identity of the compartment changes as CWP is processed and sorted (Mani & Thattai, 2016). Here, we have shown that the staging of ESVs via their morphologies demarcates ESVs undergoing unique molecular events and provides a better framework for dissecting ESV molecular biology and the developmental state of encysting cells.

**Fig. 2.**
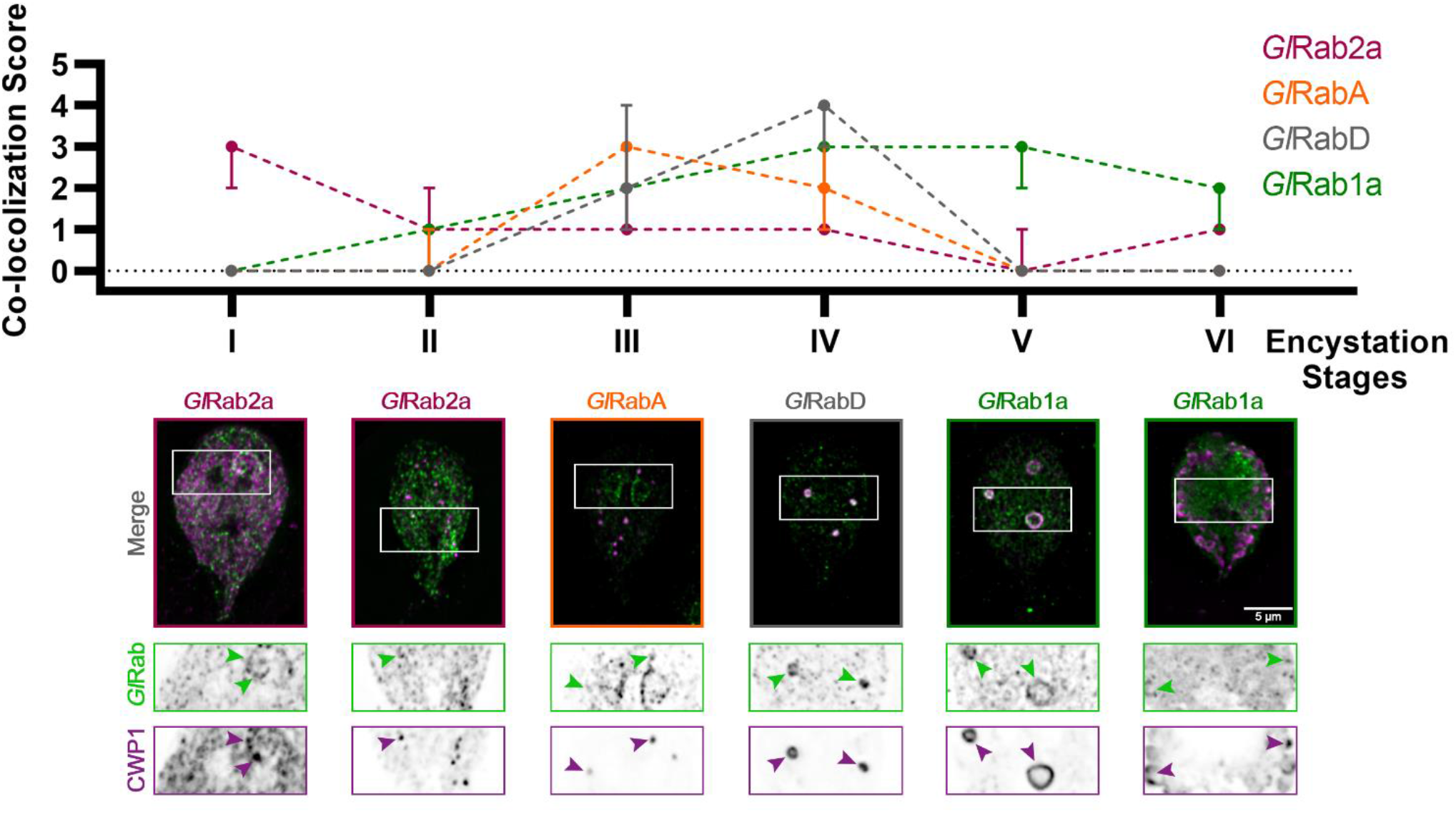
*Giardia* Rabs associate with ESVs during encystation in a stage-specific manner. Summary of findings from colocalization analysis of *Giardia* Rabs and CWP1 through all the encystation stages described above. Cells expressing endogenously tagged mNG-*Gl*Rabs were subjected to the two-step encystation process. They were then harvested at 8h and 24h p.i.e. to be fixed and stained for CWP1. 15-20 cells per encystation stage were then imaged to visualize mNG-tagged *Gl*Rabs (green) and CWP1 (magenta) and scored for the level of colocalization between the tagged *Gl*Rabs and CWP1 stained structures. Plot shows median scores with 95% confidence interval. Arrowheads indicate mNG-*Gl*Rabs colocalizing with CWP1-stained ESVs.

### *Gl*Rac activity during encystation is stage-specific

Small Rho GTPase proteins, by cycling between an active GTP-loaded conformation and an inactive GDP-bound conformation, act as molecular switches that spatially and temporally regulate the recruitment of effector proteins. We previously noted that *Giardia’s* sole Rho GTPase, *Gl*Rac, has variable association with ESVs (Krtková et al., 2016), but we did not determine the encystation stage-specificity of it. *Gl*Rac was endogenously tagged to express HALO on its N-terminus and the cells were subjected to the same ESV colocalization assay as described above. HALO-*Gl*Rac strongly colocalizes with CWP1 during Stage I, an association which wanes through the mid-stages, and then increases at Stages V and VI of encystation (Fig. 3A). This is consistent with the proposed activities of *Gl*Rac during encystation. A large proportion of *Gl*Rac is thought to be sequestered in its inactive form at the ER while *Gl*Rac association with mid-stage and late-stage ESVs promotes ESV maturation and CWP secretion, respectively (Krtková et al., 2016). We then used a CRIB domain-based Rho GTPase biosensor (Manser et al., 1994; Srinivasan et al., 2003), CRIB-mNG, to confirm our functional analyses. CRIB-mNG, which marks active GTP-loaded *Gl*Rac, colocalized with ESVs as their CWP cargo were being first sorted into condensed and fluid phases i.e. Stages III and IV, and also at Stage VI of encystation when ESVs have completed undergoing the second sorting step and were beginning to secrete their cargo to form the cyst (Fig. 3B). This pattern of localization by CRIB-mNG indicates that the ER associated population of *Gl*Rac is GDP-loaded while the *Gl*Rac associated with ESVs is GTP-loaded.

**Fig. 3.**
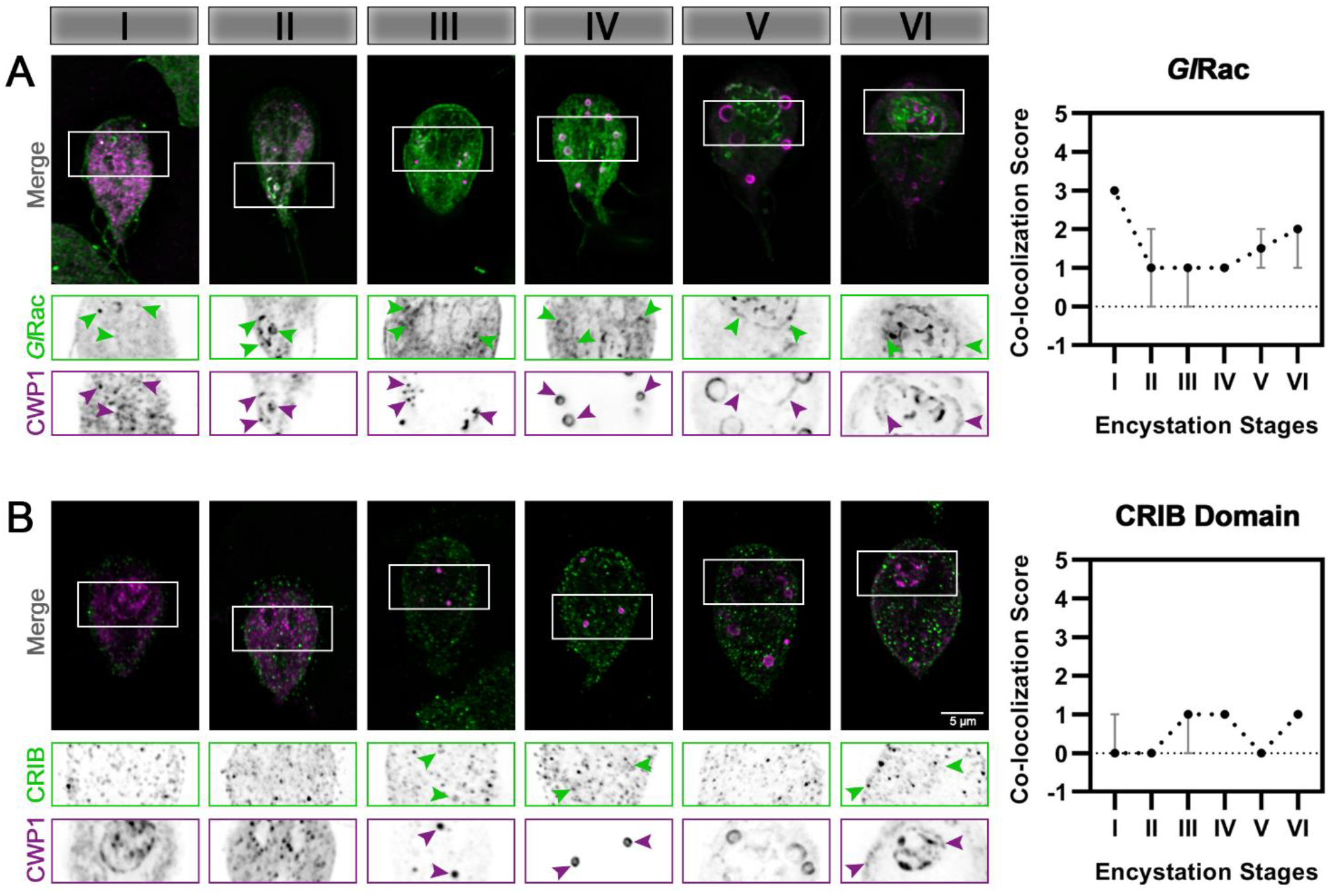
*Gl*Rac activity during encystation is stage-specific. Cells expressing HALO-*Gl*Rac (endogenous tag) or CRIB-HA-mNG (exogenous expression) were subjected to the two-step encystation process. They were then harvested at 8h and 24h p.i.e. to be fixed and stained for HALO-*Gl*Rac or CRIB-HA-mNG and CWP1. 15-20 cells per encystation stage were then imaged to visualize HALO-*Gl*Rac (green) or CRIB-HA-mNG (green) and CWP1 (magenta) and scored for the level of colocalization between HALO-*Gl*Rac or CRIB-HA-mNG and CWP1 stained structures. Plot shows median scores with 95% confidence interval. Arrowheads indicate Halo-*Gl*Rac or CRIB-HA-mNG colocalizing with CWP1-stained ESVs.

### Putative effectors of *Gl*Rac during *Giardia* encystation

Next, we sought to identify the effectors regulated by *Gl*Rac during encystation – in this study we specifically focused on mid-stage encysting cells featuring ESVs that associate with active *Gl*Rac. *Giardia* cells exogenously expressing N-terminally OneSTrEP (OS-) tagged *Gl*Rac were induced to encyst using the two-step method. We then performed an affinity purification with OS-*Gl*Rac lysates at 8 h p.i.e. Purifications were completed in triplicate, and wild type (WT) trophozoites were used as a control to identify non-specific binding to the Strep-Tactin beads. The resin eluates were then analyzed via mass spectrometry to identify putative interactors of *Gl*Rac. We identified 57 proteins, statistically significantly associating with *Gl*Rac in at least one replicate, in both trophozoites and encysting cells. Similarly, 33 proteins associated with *Gl*Rac exclusively in non-encysting trophozoites while 28 proteins were associated with encysting cells. The complete list, including low-abundance hits and proteins also identified in our mock control, is given in Table S3. Altogether, we identified 18 proteins predicted to have a role in membrane trafficking that were enriched in our encysting population (Table S4). Out of these, four were previously known to associate with ESVs supporting the idea that our list could have additional ESV components. We focused on 11 proteins that were homologs of known players of membrane trafficking in other eukaryotes (Table 1). *Gl*Rab1a and *Gl*Rab2a had already been visualized earlier (Fig. S3). Each of the rest of the candidate genes were endogenously expressed with a dual tag of triple hemagglutinin (HA) fused to mNG in cells already endogenously expressing Halo-*Gl*Rac. The candidates were then visualized via immunofluorescence assays along with *Gl*Rac and CWP1 (Fig. S4). Six of the eight candidates listed showed patterns of colocalization with CWP1 similar to that of *Gl*Rac – *Gl*Rab2a (Fig. 2), *Gl*Sec61-α, *Gl*Coatomer α subunit, *Gl*Coatomer β’ subunit and *Gl*v-SNARE (Fig. 4) suggesting that they might be involved in the same pathway regulating encystation as *Gl*Rac.

**Table 1.**
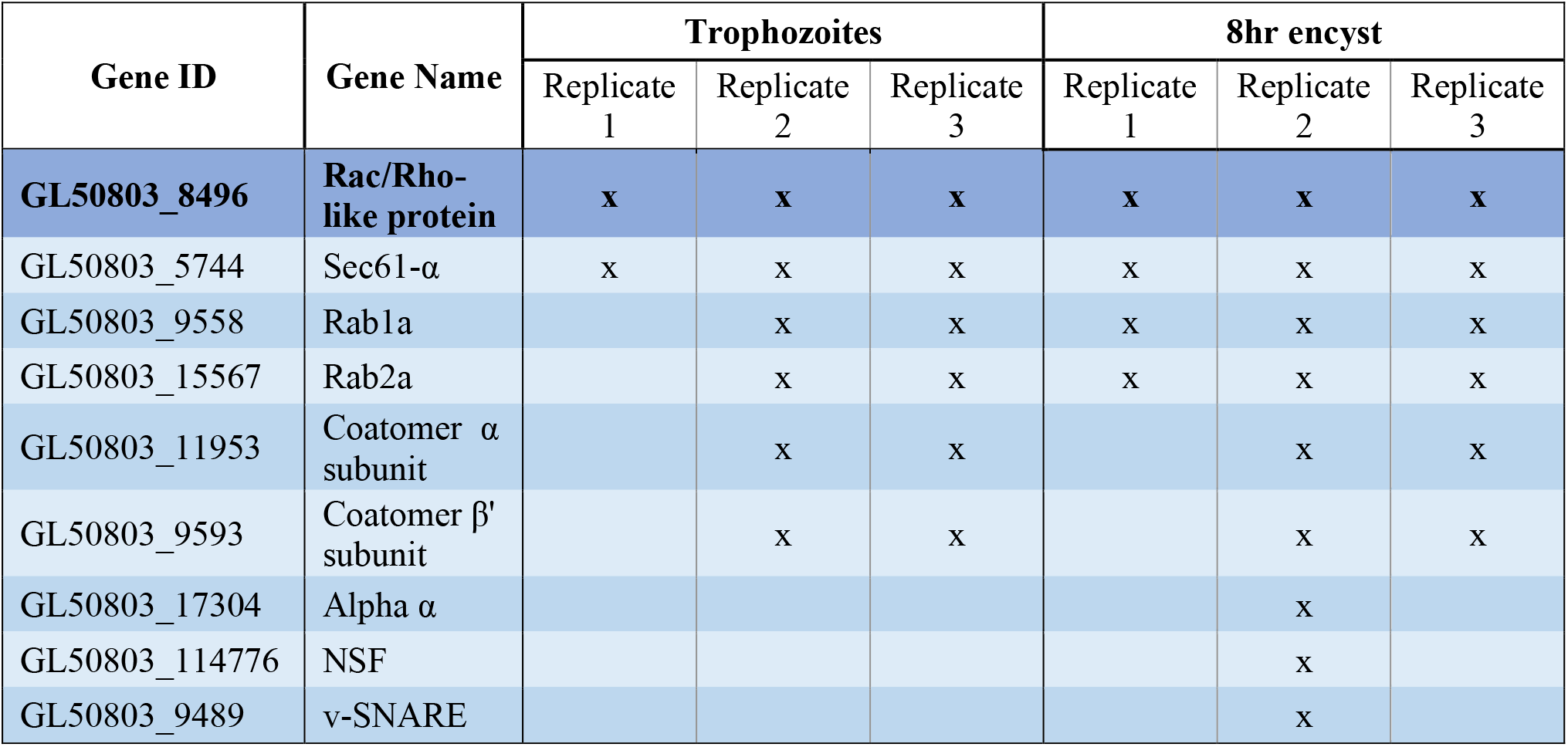
List of candidate genes selected for this study and identified as hits in OS-GlRac pulldown experiment in trophozoites and encysting cells (8h p.i.e.) that are homologs of known membrane trafficking players.

**Fig. 4.**
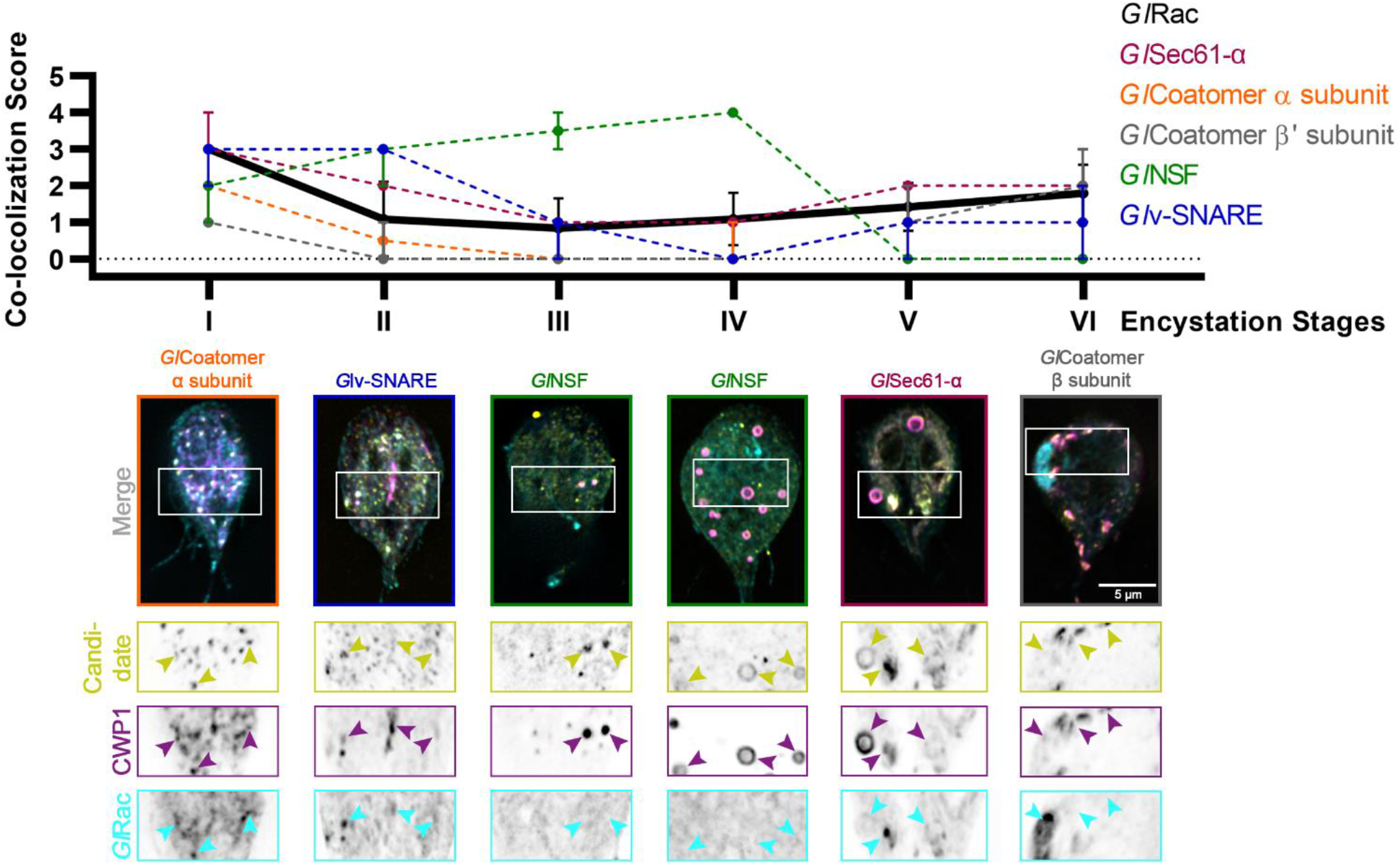
Putative effectors of *Gl*Rac colocalize with CWP1 in a stage-specific manner in a pattern that is similar to *Gl*Rac. Summary of findings from colocalization analysis of putative *Gl*Rac interactors and CWP1 through all the encystation stages described above. Cells expressing endogenously tagged HALO-*Gl*Rac and mNG-HA tagged candidates were subjected to the two-step encystation process. They were then harvested at 8h and 24h p.i.e. to be fixed and stained for CWP1. 15-20 cells per encystation stage were then imaged to visualize HALO-*Gl*Rac (cyan) or HA-mNG-candidate/candidate-HA-mNG (yellow) and CWP1 (magenta) and scored for the level of colocalization between the tagged candidate and CWP1 stained structures. Plot shows median scores with 95% confidence interval. Arrowheads indicate candidate proteins colocalizing with CWP1-stained ESVs.

## Discussion

Here, we have developed a staging system for encysting cells and have identified novel molecular markers for the specific ESV stages that correlate with the progression of encystation. We previously noted that *Gl*Rac associates with ESVs at defined points of their maturation process (Krtková et al., 2016). The lack of an established method to unambiguously pinpoint progression of encystation in individual cells, along with the variability of encystation rates that result from different *in vitro* encystation protocols, prompted us to develop a staging system that is based on ESV morphologies and independent of the method used to induce encystation. The encystation stage-specific recruitment of these molecular markers is consistent with the cisternal maturation model (Mani & Thattai, 2016) and the observations of Štefanić et al., 2009. Briefly, ESVs are thought to act as developmentally-induced Golgi (Marti et al., 2003). CWPs are first detected approximately two hours after the induction of encystation (two-step protocol); the export of CWPs from the ER, maturation of ESVs and secretion of processed CWPs to the plasma membrane together represent a simplified version of the Golgi cisternal maturation model. CWP export from the ER was found to require functional Arf1, Sar1 and Rab1 (Štefanić et al., 2009). ESVs generated *de novo* from this export process at ERESs contain pre-sorted material that is simultaneously transported and processed.

Staging ESV maturity previously relied on timing p.i.e. or simply referring to encysting cells as “early” or “late” without a standardized criteria for making these distinctions (Einarsson et al., 2016; Konrad et al., 2010; Luján et al., 1996). Timing p.i.e. is a practical approach to experimentation and can even be meaningful when comparing cells exposed to the same encystation medium when broad trends are being studied. Yet, this method is imprecise due to *in vitro* encystation protocols being only semi-synchronous. It is thought that there is a restriction point that prevents cells outside of G2 from entering the encystation response (Reiner et al., 2008). Given an eight-hour cell cycle it is not surprising that we find cells at differing encystation stages when examining induced populations. Additionally, the timing and efficiency of inducing encystation varies between the three main *in vitro* encystation methods – cholesterol starvation (Luján et al., 1996), the Uppsala method (Einarsson et al., 2016), and the two-step encystation protocol (used in this study). To a smaller extent lot-to-lot variation of serum and bile components of growth medium and encystation medium impact doubling times and, therefore, the efficiency of inducing encystation. Most importantly, *in vivo* studies employing animal models of infection currently cannot be synchronized and timing h p.i.e would not be informative. Recent efforts to study the encystation response *in vivo* has raised questions about whether *in vitro* studies can recapitulate *in vivo* encystation dynamics (Pham et al., 2017). Thus, the emerging view is that *in vitro* observations should be verified *in vivo* and this encystation staging system would also be a practical way to characterize the distribution of encysting cells within host intestines.

Our universally applicable staging system will allow the field to standardize encystation staging regardless of the method of induction used. Each ESV morphology change is thought to correspond with sequential molecular events, including secretion from the ER, CWP processing, sorting, and pulsed cellular secretion of processed CWP. ESV morphology changes therefore provide landmarks for encystation stages. Observing and categorizing ESV morphologies are obvious and easy to follow, therefore making it accessible for adoption by the wider *Giardia* research community. This would be advantageous since it would reduce ambiguity when the timeline of molecular pathways directing CWP traffic are being specified. Finally, as the resolution of these events increase, encystation stages could be further subdivided to accommodate new discoveries.

Beyond morphological categorization we have identified several *Giardia* Rab GTPases that demarcate specific encystation stages, indicating that our staging system corresponds to unique molecular identities. Some Giardia Rab GTPases were previously known to be associated with ESVs and overall this group of proteins is upregulated during encystation (Einarsson et al., 2016; Marti et al., 2003). Here we tagged and followed eight out of nine *Giardia* Rab GTPases over the course of encystation. Seven of the eight Rab GTPases analyzed here associated with ESVs. By scoring the degree of colocalization with CWP1 during the different encystation stages, we were able to identify their distinct patterns of ESV association thus potentially indicating the point at which their functions were required as encystation progressed. Functional equivalence between some of the *Gl*Rabs and their homologs can be inferred. *Gl*Rab2a and CWP1 colocalization peaked at Stages I and II, paralleling its mammalian homologs which reside in the ER-to-Golgi Intermediate Compartments (ERGIC) and regulate Golgi biogenesis, and bidirectional transport between the two compartments (Saraste, 2016). *Gl*RabD shows the closest homology to Rab8 or Rab13 known to localize at the *trans* Golgi network, recycling endosomes, late endosomes and the plasma membrane. They have especially been implicated in biosynthetic and recycling endosomal pathways (Ioannou & McPherson, 2016). In *G. lamblia*, RabD colocalization with ESVs peaks at Stage IV and equivalence could be argued.

While the similarities between *Giardia* Rabs and their eukaryotic homologs allow for some inferred function, not all *Giardia* Rabs have clear orthologs. The closest homologs of *Gl*RabA (RabA in plants/Rab 4 or Rab11 in mammals) are known to associate with the *trans* Golgi and post-Golgi networks, specifically, the recycling endosomes (Li & Marlin, 2015; Minamino & Ueda, 2019; Vernoud et al., 2003) and *Gl*RabA was shown to associate with ESVs the most at Stage III, which is likely equivalent to *cis* Golgi earlier in canonical secretory pathway. Additionally, *Gl*Rab1a colocalization with ESVs peaks at Stages V and VI which is when the ESVs undergo further sorting and the fluid phase of ESVs are beginning to disassociate for secretion. This is different from canonical Rab1 isoforms that have been established to regulate transport between ER-ERGIC-Golgi interface (Saraste, 2016).

The importance of Rabs in regulating *Giardia* encystation is yet to be fully understood. Previously, *Gl*Rab1a was shown to be necessary for ESV development and cyst wall formation (Štefanić et al., 2009) complementing the observations we have made here. The specific functions of *Gl*Rabs would make an interesting topic for future studies.

As a proof of principle for our staging system, we turned to *Gl*Rac which has a complex relationship to ESVs. In agreement with previously published data, *Gl*Rac colocalization with CWP1 peaked at stages I and VI with low level colocalization being detectable throughout the rest of the stages. *Gl*Rac was hypothesized to be sequestered in the ER in an inactive state and then have a role in promoting ESV maturation and secretion of CWP1 (Krtková et al., 2016). Based on our hypothesis, *Gl*Rac was expected to be active at Stages III/IV and VI which we confirmed using CRIB-mNG as a *Gl*Rac signaling biosensor. As a molecular switch, *Gl*Rac is expected to recruit effectors that promote maturation of ESVs and secretion of CWP1.

To identify *Gl*Rac effector proteins we affinity purified *Gl*Rac interactors from non-encysting and encysting populations. We were not surprised to find many statistically significant interacting proteins since *Gl*Rac, as the sole Rho GTPase in *Giardia*, is presumably responsible many of the same roles the multitude of Rho GTPases carry out in other eukaryotes (Hodge & Ridley, 2016; Lawson & Ridley, 2018; Phuyal & Farhan, 2019). Here we chose to focus on proteins predicted to have a role in trafficking as we had the potential to identify novel ESV components. Indeed, every protein selected from the proteomics hits associated with the ER and/or ESVs with *Gl*Rac.

A number of candidates displayed patterns of colocalization that were similar to the patterns displayed by *Gl*Rac with CWP1, including *Gl*Sec61-α, *Gl*Coatomer-α subunit, *Gl*Coatomer-β’ subunit and *Gl*v-SNARE. Each of these proteins was found to be in the ER at Stage I with high colocalization scores with CWP1 which reduced in subsequent stages to rise back up again in the final two stages. This would suggest that they localize in the same compartments during encystation. Effector proteins can also bind the inactivated GDP-bound form of Rho GTPases, so it is not surprising that some of these candidates colocalize with *Gl*Rac at the ER where *Gl*Rac is largely GDP-loaded. All of these except *Gl*v-SNARE were also pulled down by *Gl*Rac in trophozoites indicating additional constitutive roles, while *Gl*NSF, *Gl*α-adaptin and *Gl*v-SNARE appears to have a encystation-specific association with *Gl*Rac, which will require further investigation. Here, we have set the scene for future studies to dissect the role of these newly identified components.

In summary we have devised a universally applicable ESV staging system based on ESV morphology using CWP1 as a marker. As CWP1 is an easily accessible encystation marker, our staging system can be readily adopted across the field. Beyond morphological differences in ESV stages we have identified molecular markers that can be used to distinguish different stages. As the *Giardia* toolkit grows, the field will have greater access to tools that allow for functional studies. We anticipate future studies of *Gl*Rabs and other ESV-associated proteins identified here that will impact ESV maturation and ESV morphology. Therefore, morphology alone may be insufficient for characterizing/staging the resulting ESVs. The newly identified molecular markers for different ESV stages will be a powerful tool for the purpose of characterizing the stages of abnormal ESVs.

## Author Contributions

ET and AP designed the experiments; ET and RS performed the colocalization with CWP1 analyses; ET and JK performed the affinity purifications. HS performed expression analysis. RJ performed mass spectrometry analysis and analysed the proteomics data. ET and AP wrote the manuscript. MM and AP were responsible for fund acquisition. All authors contributed to the article and approved the submitted version.

## Acknowledgments

This work was supported by NIH Grant 5R01AI110708 (A.R.P.) and funding provided by the NIH Yeast Resource Center P41 GM103533 (M.M) supported generation of the mass spectrometry data in part. Authors acknowledge Kelli L. Hvorecny and Melissa Steele-Ogus for their input and critical reading of the manuscript.

---

**Fig. S1.**
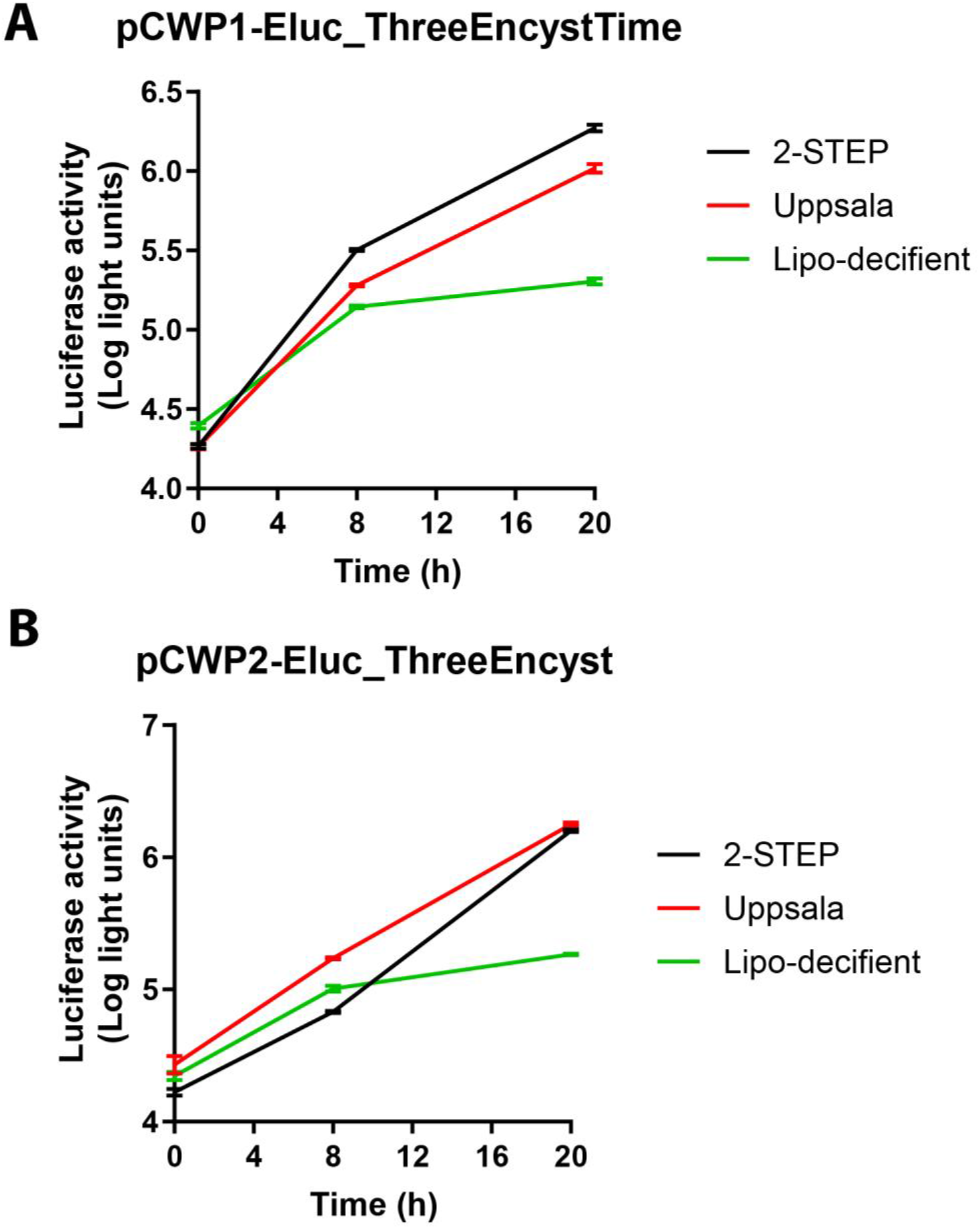
Comparison of CWPl-Ehic and CWP2-Ekic reporter strains using three common methods to induce encystation in Giardia. This is a single experiment with three separate technical replicates for each condition and timepoint. While the CWP1 result for Two-Step and Uppsala appear similar in this bulk assay, it should be noted that the high bile levels used in the Uppsala method kill off a portion of the parasites and this results in a stronger induction of encystation for the surviving cells.

**Fig. S2.**
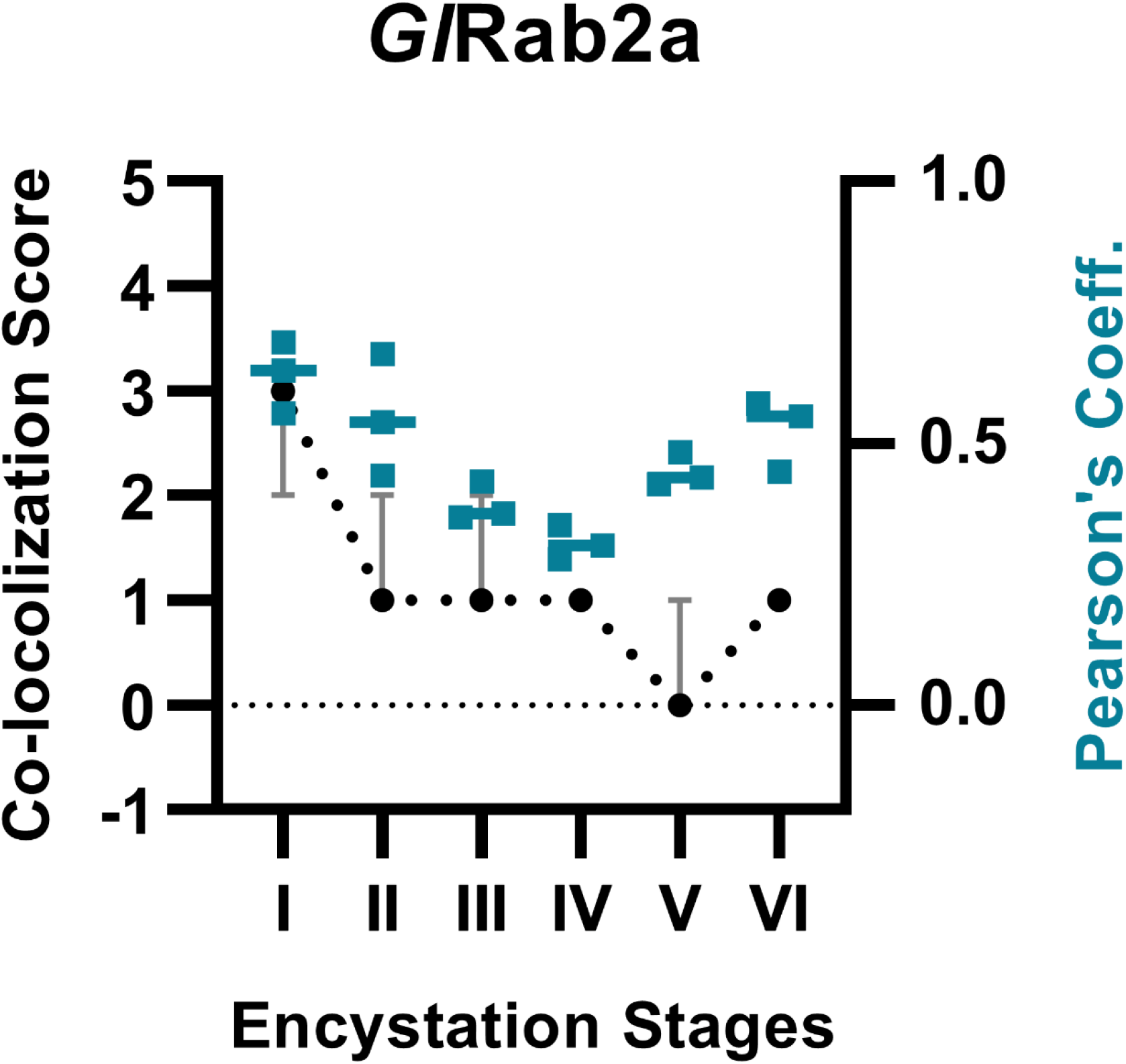
Comparing manual scores and Pearson’s MAS Correlation Co-efficients for colocalization levels between *GI*Rab2a and CWP1 across the 6 encystation stages.Plot shows median scores with 95% confidence interval.

**Fig. S3.**
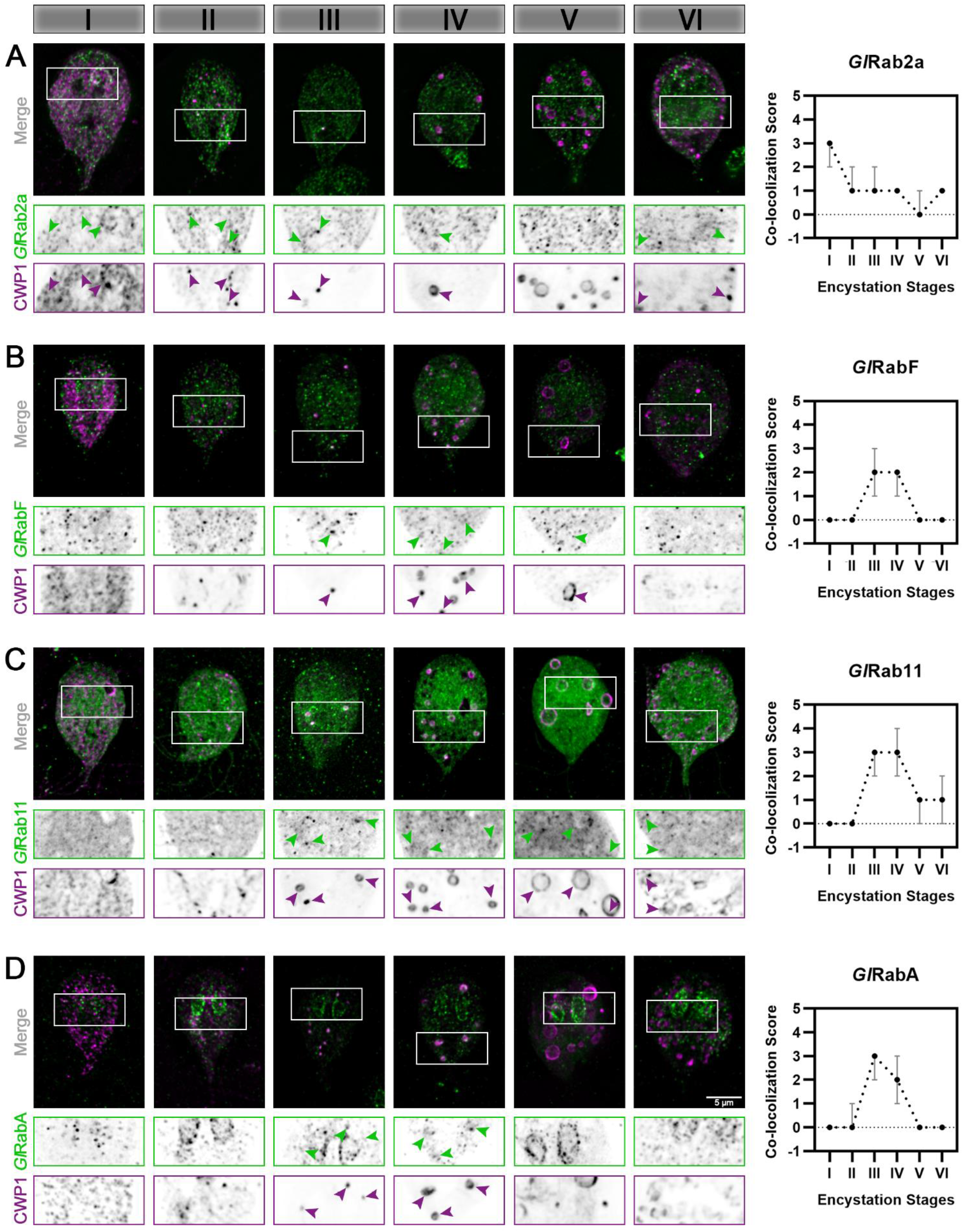
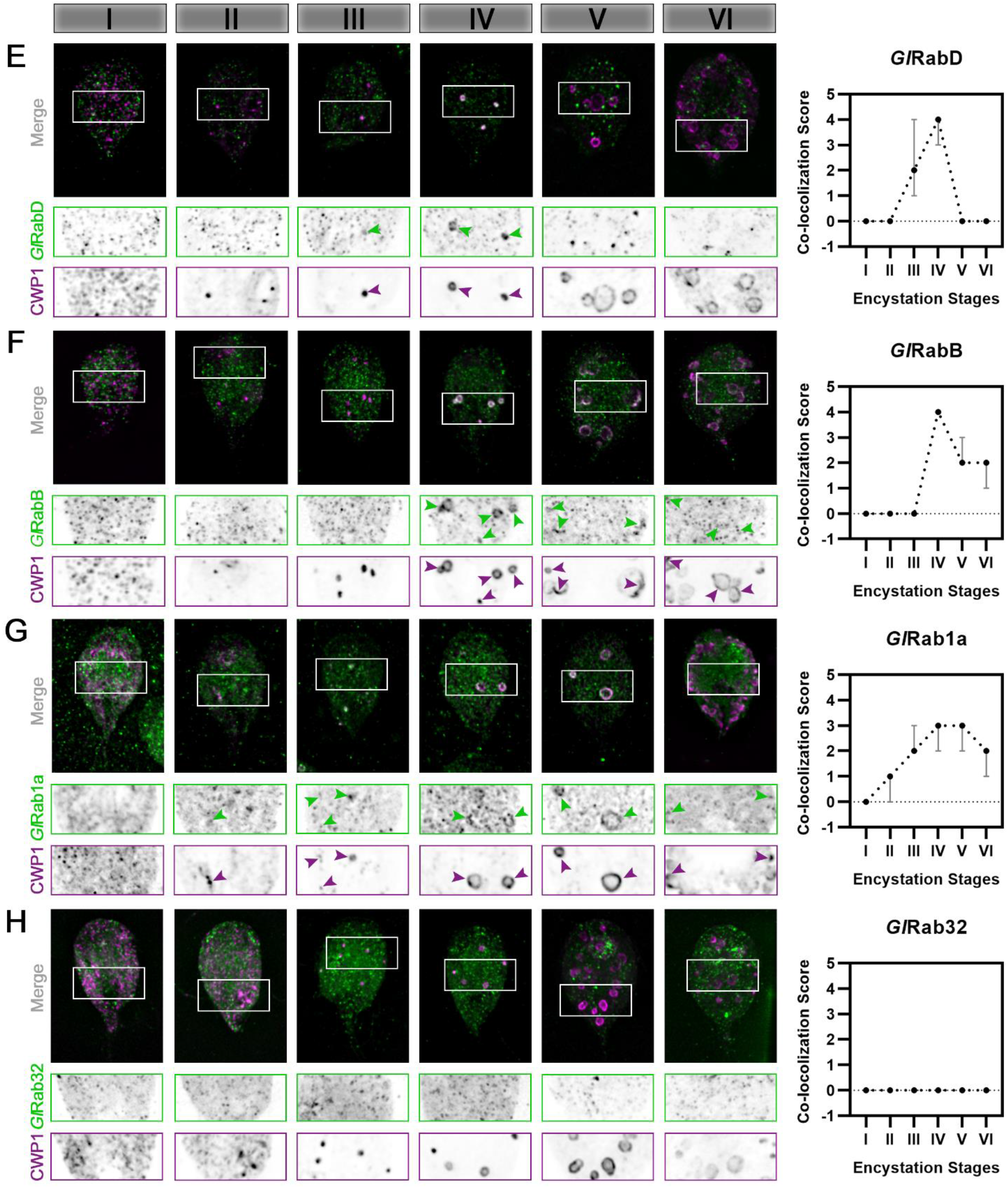
*Giardia* Rabs associate with ESVs during encystation in a stage-specific manner. Colocalization analysis of *Giardia Rabs* and CWPI through the encystation stages. Cells expressing endogenously tagged mNG-*GI*Rabs were subjected to the two-step encystation process. They were then harvested at 8 h and 24 h p i e to be fixed and stained for CWP1. 15-20 cells per encystation stage were then imaged to visualize mNG-tagged *GI*Rabs (green) and CWP1 (magenta) and scored for the level of colocalization between the tagged *GI*Rabs and CWP1 stained structures. Plots show median scores with 95% confidence interval. Arrowheads indicate mNG-*GI*Rabs colocalizing with CWPI-stained ESVs.

**Fig. S4.**
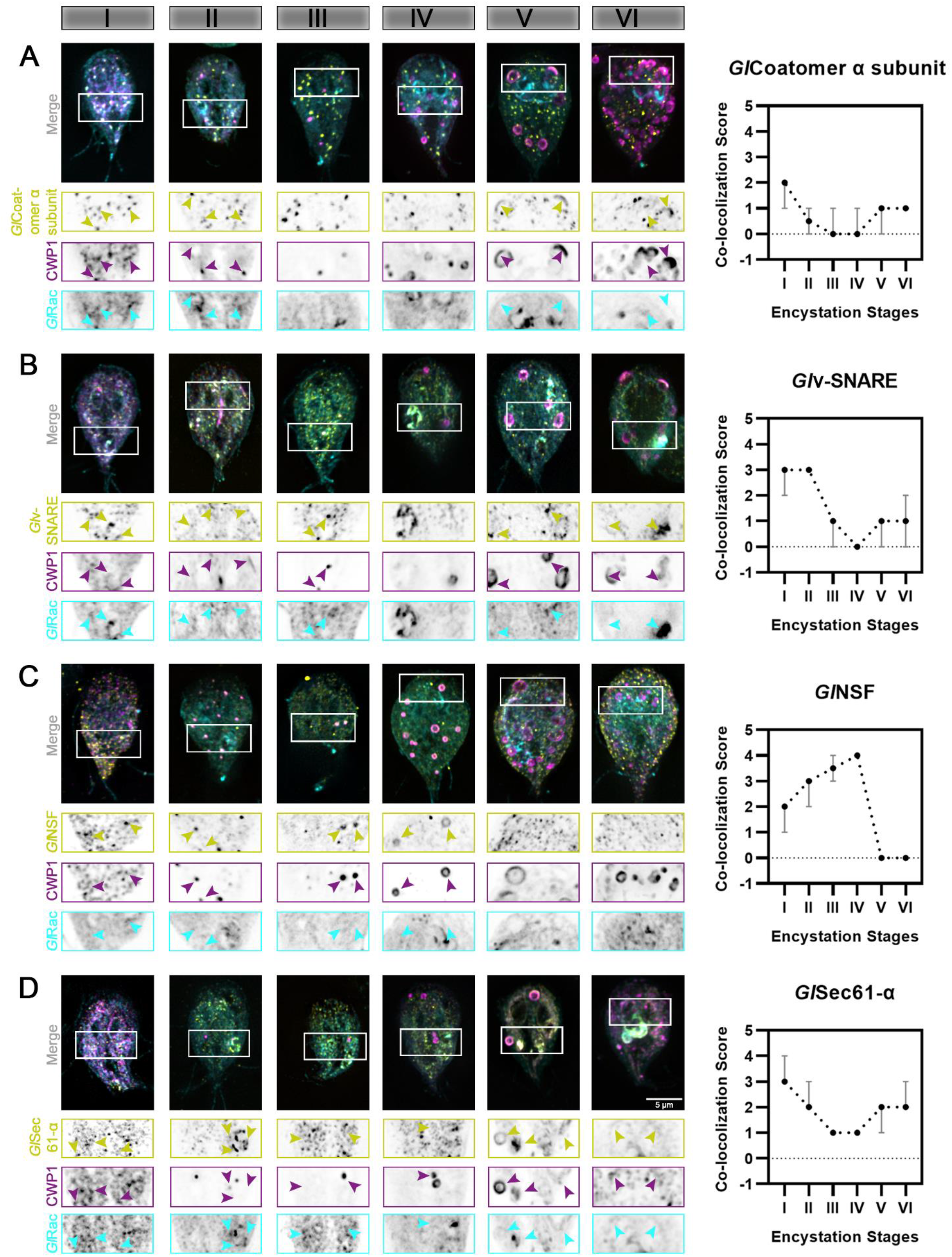
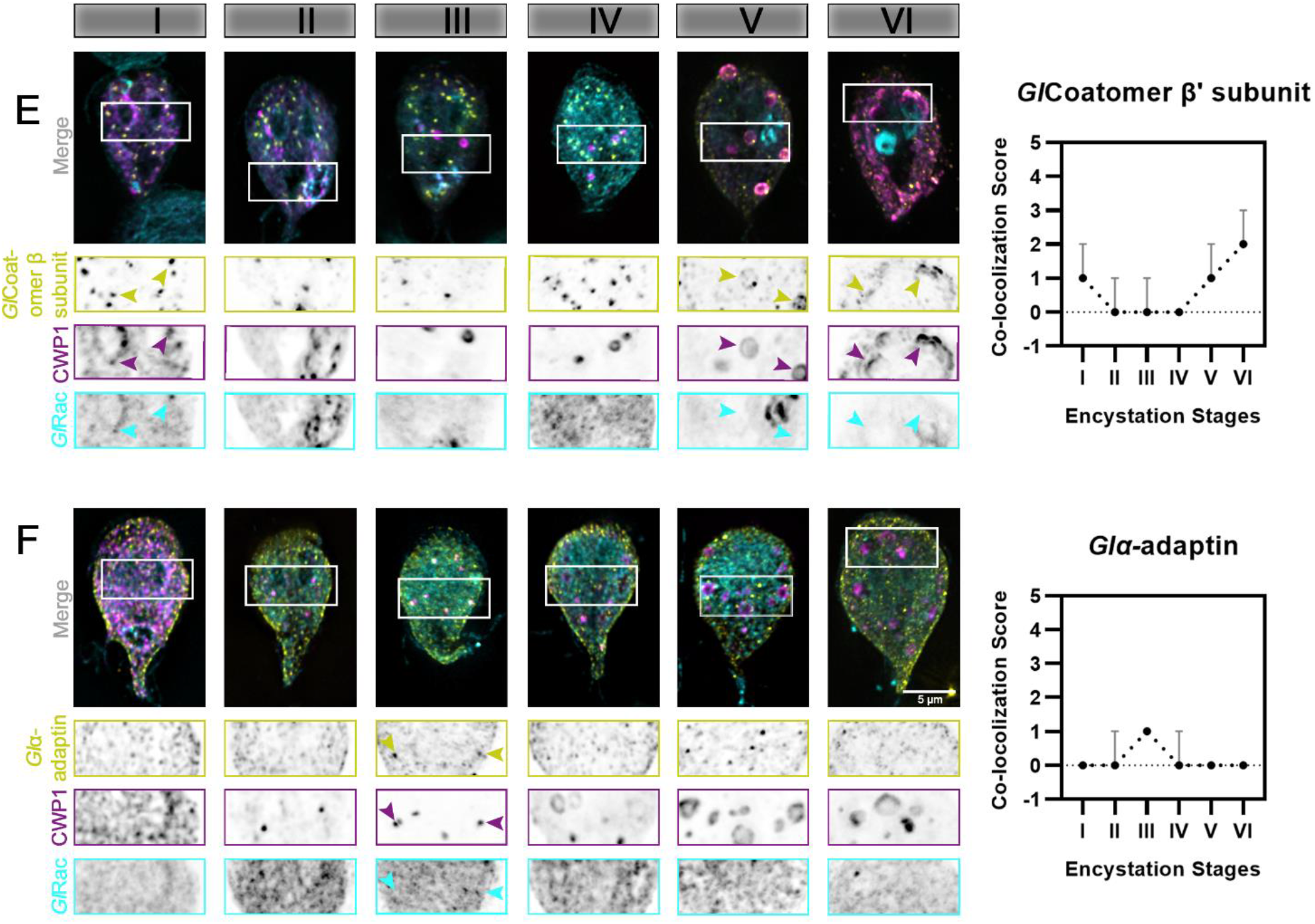
Putative effectors of *GI*Rac colocalize with CWPI in a stage-specific manner. Colocalization analysis of putative *GI*Rac interactors and CWPI through all the encystation stages described above. Cells expressing endogenously tagged HALO-*GI*Rac and mNG-H A tagged candidates were subjected to the two-step encystation process. They were then harvested at 8 h and 24 h p.i.e. to be fixed and stained for CWPI. 15-20 cells per encystation stage were then imaged to visualize HALO-*GI*Rac (cyan) or HA-mNG-candidate/ candidate-HA-mNG (yellow) and CWP1 (magenta) and scored for the level of colocalization between the tagged candidate and CWP1 stained structures. Plot shows median scores with 95% confidence interval. Arrowheads indicate candidates colocalizing with CWPI-stained ESVs.

**Fig. S5.**
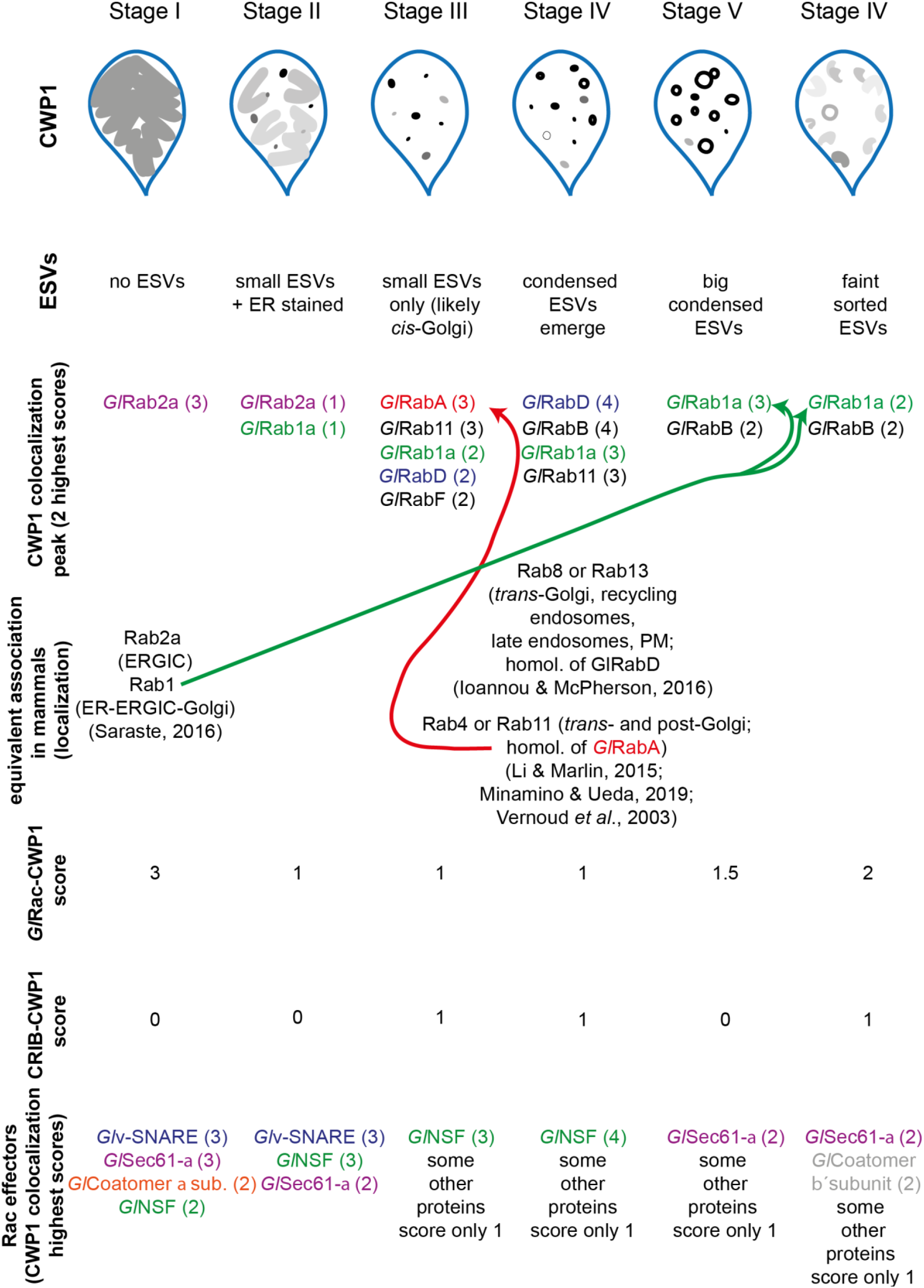
Graphical summary showing the relationship of all proteins studied here relative to ESV stages.

